# Engineered probiotic *Escherichia coli*-mediated intestinal nicotine clearance alleviates nonalcoholic steatohepatitis in mice

**DOI:** 10.64898/2026.07.02.736048

**Authors:** Ni Zuo, Xin Cai, Weitong Wang, Zequan Ren, Zihan Jiang, Weihong Jiang, Xin Song, Yang Gu

## Abstract

Nicotine accumulates in the gut and drives non-alcoholic steatohepatitis (NASH) via the gut-liver axis, yet no effective clinical intervention is currently available. To address this challenge, the probiotic *Escherichia coli* Nissle 1917 (EcN) was engineered for in situ nicotine clearance in the gut. Mutational screening of nicotine oxidoreductase 2 (PpNicA2) identified a highly active variant, PpNicA2^A107R^. Its incorporation into EcN together with an electron transfer protein (CycN) and a newly identified transporter (T3/T7) yielded 80% nicotine-degrading activity. Chromosomal integration of this module generated a stable strain, EcN-N12, which in NASH mouse models depleted intestinal nicotine, rescued hepatic lipid metabolism, alleviated tissue damage, and intercepted the nicotine-mediated gut-liver axis pathological progression. This work thus offers an effective and clinically translatable approach for nicotine-associated diseases.

## Introduction

Nicotine is a toxic, amphipathic alkaloid primarily derived from *Nicotiana* species^1^. It can readily cross biological membranes, entering the body via the respiratory tract, skin, and intestines, where it stimulates the nervous and cardiovascular systems, increasing heart rate, blood pressure, and blood viscosity^2^. Excessive nicotine exposure elicits adverse reactions including nausea and vomiting, with lethal consequences observed under extreme overdose^3^. Chronic nicotine ingestion further impairs gastrointestinal^4^ and neurological functions^5^, causes multi-organ lesions^6^, and disturbs hepatic lipid homeostasis^7^. Recent studies reveal significant nicotine accumulation in the intestinal tissues of smokers^8^, and animal experiments indicate that high ileal nicotine levels promote intestinal ceramide production and release^9^, exacerbating hepatic steatosis, inflammation, and fibrosis, thus driving the progression of non-alcoholic fatty liver disease (NAFLD)^10^. Given the FDA approval and future promotion of oral nicotine pouches, the risk of gut nicotine-induced diseases will likely increase^11–13^, highlighting the urgent need for safe, effective interventions to clear intestinal nicotine and reduce disease risk.

Physical and chemical nicotine degradation methods, e.g., adsorption, solvent extraction, chemical catalysis^14, 15^, are not applicable in the human body. Considering the intestinal microenvironment, leveraging gut microbial metabolism to eliminate nicotine is a feasible approach. To date, three microbial nicotine degradation pathways have been identified, i.e., the pyridine pathway (predominantly found in *Arthrobacter* spp.)^16^, the pyrrolidine pathway (predominantly found in *Pseudomonas* spp.)^17–19^, and a hybrid of the pyridine and pyrrolidine (HPP) pathway (primarily found in *Agrobacterium tumefaciens*)^20, 21^. Nicotine dehydrogenase (NdhAB) catalyzes the first step in both the pyridine and HPP pathways, converting nicotine to 6-hydroxynicotine ^21^, which then undergoes a six-step metabolic cascade to yield 2,3,6-trihydroxypyridine and succinate. In contrast, the pyrrolidine pathway, driven mainly by nicotine oxidoreductase 2 (PpNicA2), is more direct: it converts nicotine to non-toxic, non-addictive *N*-methylmyosmine^22–24^, which is then hydrolyzed to pseudooxynicotine and sequentially oxidized to 3-succinoylpyridine, 6-hydroxy-3-succinoylpyridine, and finally to fumaric acid, which enters central carbon metabolism.

While the catabolic routes of nicotine have been characterized, the identity of nicotine transporters in bacteria remains elusive. This knowledge gap is critical, as the transmembrane transport of nicotine into bacterial cells constitutes a rate-limiting step in the entire degradation pathway^25, 26^. Efficient transporters not only enhance substrate uptake efficiency but also ensure the sustained flux of intracellular metabolism, thereby substantially improving overall degradation capacity. Hence, the identification of functional nicotine transporters represents an essential step toward the rational engineering of highly efficient nicotine-degrading bacteria.

In this study, the probiotic *E. coli* Nissle 1917 (EcN), a safe, genetically tractable, and gut-adaptive strain^27–29^, was selected as the chassis for engineering an efficient nicotine-degrading bacterium. A highly active dehydrogenase for nicotine oxidation was identified and its capacity was further augmented through co-expression with an electron acceptor. In addition, two efficient nicotine transporters, designated T3 and T7, were discovered. The engineered EcN strain harboring the composite module (PpNicA^A107R^-CycN-T7-T3) achieved efficient nicotine degradation in vitro and in vivo. In animal studies, this strain effectively alleviated nicotine-exacerbated non-alcoholic steatohepatitis (NASH) injury and demonstrated favorable biosafety.

## Results

### Assembly of an efficient nicotine bioconversion pathway

The first dehydrogenase identified to oxidize nicotine was PpNicA2 (hereafter PpNicA), an FAD-dependent flavin dehydrogenase from *Pseudomonas putida*^30^. Its variants, PpNicA^A107R^ and PpNicA^F355H24^, showed 19-fold and 13.4-fold, respectively, higher activity than the wild-type^31^. Recently, the enzyme BxNicX from *Bacteroides xylanisolvens* was found to oxidize nicotine to 4-hydroxy-1-(3-pyridyl)-1-butanone (HPB)^9^. Domain analysis and sequence alignment of PpNicA, its variants (A107R and F355H), and BxNicX (Fig. 1a, Supplementary Fig. 1) reveal marked structural differences: PpNicA and its variants share a conserved FAD-dependent oxidoreductase core, while BxNicX exhibits a distinct RVT1/GIIM-dominated architecture, pointing to significant divergence between the two enzymes.

**Fig. 1.**
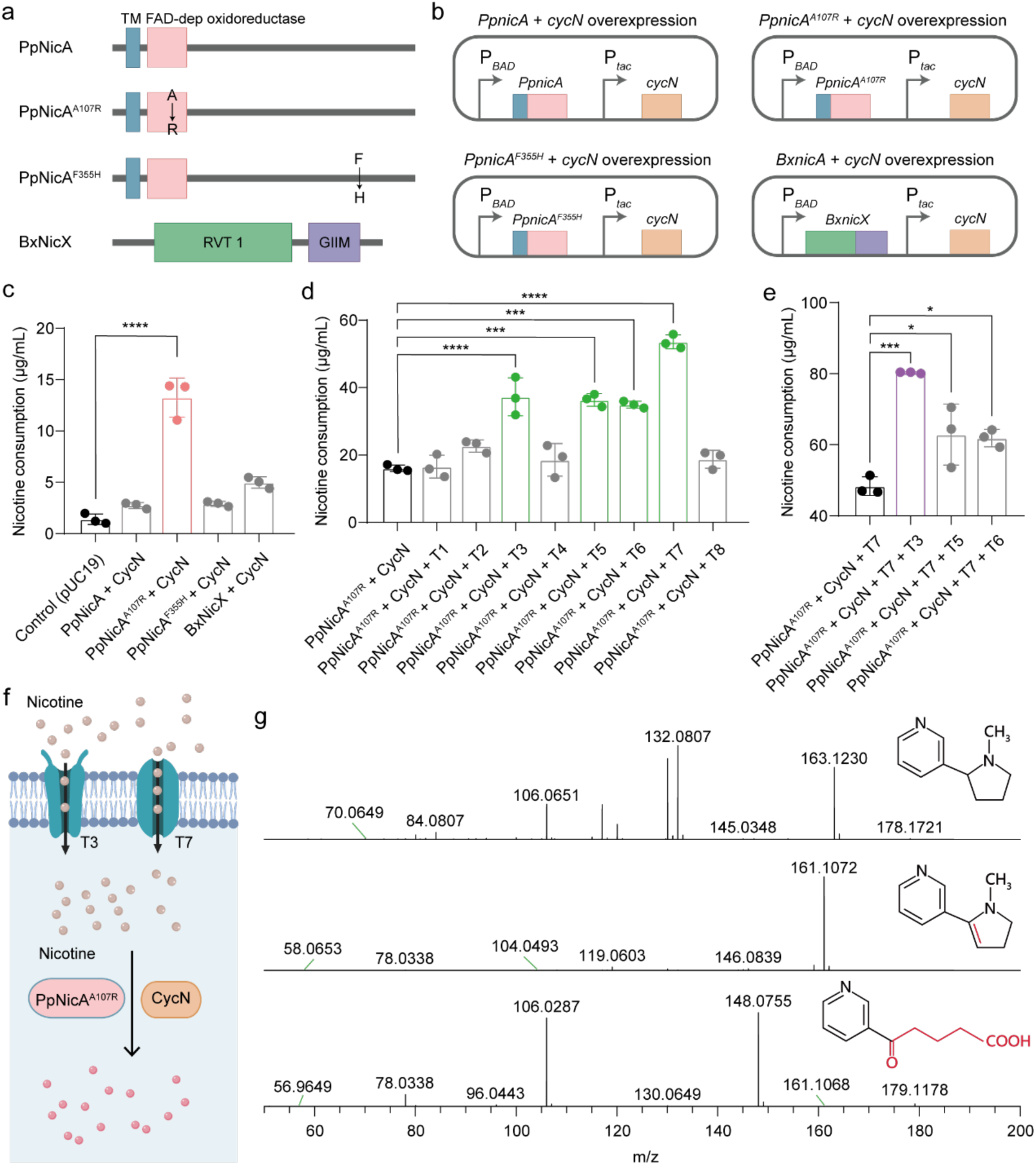
Screening of efficient nicotine-degrading elements. a, Schematic diagrams of the protein structures of NicA family members from different sources. b, Schematic illustration of the co-expression of *nicA* family genes with *cycN* in EcN. c, HPLC analysis of the degradation of 100 μg/mL nicotine by EcN strains co-expressing *nicA* family genes and *cycN*. d, HPLC analysis of nicotine (100 μg/mL) degradation by EcN strains co-expressing *PpnicA^A^*^107^*^R^*-*cycN* with each of eight candidate nicotine transporters genes *t1*-*t8*. Statistical analysis was performed using one-way ANOVA with Tukey’s multiple comparison test. \**P* < 0.05, \*\*\**P* < 0.001, \*\*\*\**P* < 0.0001. e, HPLC verification of nicotine (100 μg/mL) degradation by EcN strains co-expressing *PpnicA^A^*^107^*^R^*-*cycN* and combinations of *t7* with *t3*, *t5*, or *t6*. f, Schematic overview of the de novo construction of PpNicA^A107R^-CycN-T7-T3 metabolic pathway in EcN. g, Qualitative detection and structural analysis of nicotine metabolites by liquid chromatography–mass spectrometry (LC-MS).

Given these structural differences, which may translate into distinct catalytic behaviors in the *E. coli* host, we next sought to systematically evaluate the nicotine-oxidizing performance of each enzyme in EcN. To this end, the coding genes of PpnicA, PpnicA^A107R^, PpnicA^F355H^, and BxnicX were individually overexpressed in EcN under the control of an arabinose-inducible promoter P*_BAD_*. In parallel, the electron acceptor CycN, under the constitutive promoter P*_tac_*, was co-expressed with each dehydrogenase, generating four expression plasmids (Fig. 1b). Each construct was then transformed into EcN and evaluated for nicotine conversion efficiency. Among the four strains, the one co-expressing the genes coding for PpnicA^A107R^ and CycN exhibited the highest nicotine conversion efficiency, converting approximately 13% of nicotine within 2 h of incubation (Fig. 1c). In contrast, the strain expressing PpnicA^A107R^ alone showed a markedly lower efficiency, with a reduction of about 59% (Supplementary Fig. 2a, b). Supplementation with exogenous FAD and FMN, the flavin-dependent dehydrogenase cofactors, did not further improve conversion (Supplementary Fig. 3), suggesting that endogenous levels of these cofactors in EcN are already sufficient to sustain PpnicA^A107R^ activity. The other three constructs, PpnicA-CycN, PpnicA^F355H^-CycN, and BxnicX-CycN, achieved conversion rates of approximately 2.7%, 2.9%, and 4.9%, respectively (Fig. 1c).

To further enhance nicotine conversion efficiency, we next turned our attention to nicotine transport into cells. Although no nicotine-specific transporter has been reported in bacteria to date, a previous proteomic study revealed that eight membrane proteins in *Pseudomonas putida* were significantly upregulated under nicotine-induced conditions^32^, indicating their potential involvement in nicotine uptake. These candidates (T1‒T8) (Supplementary Table 1) were therefore tested in the PpnicA^A107R^-CycN-harboring EcN strain described above. Encouragingly, introduction of T3, T5, T6, or T7 each enhanced nicotine degradation in the engineered EcN strain, with T7 proving the most effective, improving conversion efficiency by approximately 54% (Fig. 1d). Co-overexpression of T7 with T3, T5, or T6 further boosted the efficiency relative to T7 alone, and the T7-T3 pair achieved the highest level, exceeding 80% conversion (Fig. 1e). Based on these results, a functional module comprising PpNicA^A107R^, CycN, T3, and T7 was assembled (Fig. 1f), and the resulting strain was designated EcN-P4. Metabolic profiling revealed that EcN-P4 converted nicotine to *N*-methylmyosmine, a non-toxic, non-addictive metabolite, which subsequently underwent spontaneous hydrolysis to pseudooxynicotine (Fig. 1g).

### Construction of a nicotine-degrading EcN strain via chromosomal integration

To circumvent the potential plasmid instability in the host strain, we sought to integrate the module PpNicA^A107R^-CycN-T7-T3 into the chromosome of EcN. Briefly, multiple previously reported neutral integration sites ^33–35^ and nonessential genes^36–41^ in EcN were selected as insertion targets (Fig. 2a). The P*_BAD_*-PpNicA^A107R^-CycN (using the promoter P*_BAD_*) cassette was sequentially inserted into the *ldhA*, *slmA*, *xylA*, *aslA*, *glmS*, *ksgA*, *kefB*, *maeB*, *nth*, *tkrA*, and *yjcS* loci via CRISPR/Cas9-mediated genome editing, yielding strains harboring 1, 3, 5, 7, 9, or 11 copies (in that order). In parallel, the P*_sal_*-T7-T3 fragment was inserted at the *rhtB* locus (a reported neutral integration site)^35^ (Fig. 2a). The resulting strains were designated EcN-N2 (1 copy), EcN-N4 (3 copies), EcN-N6 (5 copies), EcN-N8 (7 copies), EcN-N10 (9 copies), and EcN-N12 (11 copies). We then evaluated the nicotine conversion capacity of these strains. EcN-N2 showed negligible activity (Fig. 2b), whereas EcN-N4, EcN-N6, EcN-N8, EcN-N10, and EcN-N12 exhibited progressively increasing conversion rates of 28%, 45%, 79%, 84%, and 89%, respectively (Fig. 2b). Notably, EcN-N12 achieved the highest efficiency, comparable to that of the plasmid-based strain EcN-P4 (Fig. 2b).

**Fig. 2.**
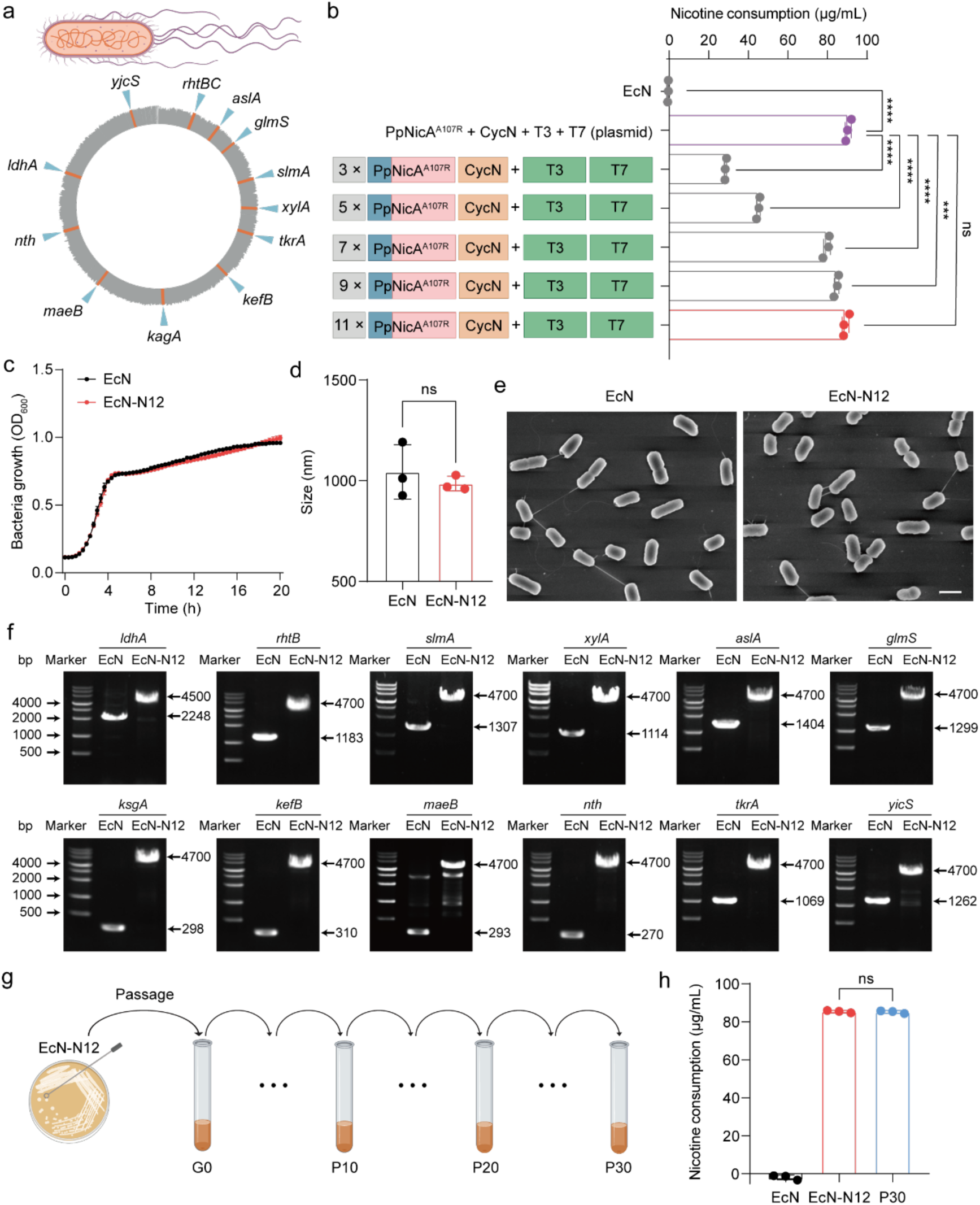
Construction and characterization of engineered probiotics with genomically integrated degradation elements. a, Schematic diagram of the 12 neutral genomic loci selected in EcN genome. b, HPLC analysis of nicotine (100 μg/mL) degradation by EcN strains with different copy numbers of degradation elements integrated into the genome. Statistical analysis was performed using one-way ANOVA with Tukey’s multiple comparison test. ns, no significance. c, Growth curves of the engineered probiotic EcN-N12 and wild-type EcN. d, Measurement of the cell size of EcN-N12 and EcN. Statistical analysis was performed using an unpaired two-tailed Student’s t-test. \*\*\**P* < 0.001, \*\*\*\**P* < 0.0001. e, SEM images of EcN-N12 and EcN. Scale bar, 1 μm. f, PCR gel verification of the sizes of the inserted fragments at each genomic editing locus in EcN-N12, with EcN as the control. g, Schematic illustration of 30 serial antibiotic-free passages of EcN-N12. h, HPLC analysis of nicotine (100 μg/mL) degradation by EcN-N12 and the passaged strain EcN-N12-P30. Statistical analysis was performed using one-way ANOVA with Tukey’s multiple comparison test. ns, no significance.

We next systematically characterized the key performance attributes of EcN-N12. The strain exhibited a growth curve similar to that of wild-type EcN (Fig. 2c), and no significant differences in cell size or morphology were observed (Fig. 2d, e), suggesting that sequential chromosomal integration of exogenous genes imposed no notable metabolic burden. Genetic stability of the integrated module (PpNicA^A107R^-CycN-T7-T3) was then assessed by serial passaging EcN-N12 for 30 generations without antibiotic selection, with colony purity monitored by streaking every 10 generations (Fig. 2g). The resulting population, designated EcN-N12-P30, was subjected to both genotypic and functional characterization. PCR analysis confirmed intact retention of all 12 integrated fragments in the genome (Fig. 2f), and the nicotine-degrading capacity of EcN-N12-P30 remained comparable to that of the parental strain (Fig. 2h). Collectively, these results demonstrate that EcN-N12 possesses robust genetic stability and sustained nicotine-metabolic activity.

### EcN-N12 ameliorates nicotine-exacerbated nonalcoholic steatohepatitis in Mice

Nicotine accumulation in humans is associated with multiple diseases^42–44^. While EcN-N12 exhibits potent nicotine-degrading activity *in vitro*, its capacity to mitigate nicotine-induced pathology *in vivo* has not been tested. Here, we focused on nonalcoholic steatohepatitis (NASH), a disease exacerbated by nicotine via the gut-liver axis^9^, and established a murine NASH model by administering nicotine in drinking water combined with a high-fat and high-sugar diet to evaluate whether EcN-N12-mediated nicotine degradation can alleviate NASH pathology *in vivo*.

A total of seven mouse groups were established in this study (Fig. 3a). Among them, mice in the nicotine + HFHCD + EcN-N12 group (hereafter referred to as the “EcN-N12 treatment group”) exhibited the slowest body weight gain, comparable to that of the healthy group, whereas the other groups, particularly the model group and the HFHCD-only group, showed pronounced increases in body weight (Fig. 3b). Consistently, the EcN-N12 treatment group had the lowest liver-to-body weight ratio (Fig. 3c), indicating that EcN-N12 ameliorated body weight gain and reduced relative liver weight induced by nicotine combined with a high-sugar high-fat diet. Consistent with these findings, hepatic and serum biochemical parameters (including TG, TC, FFA, AST, ALT, IL-6, and TNF-α) revealed that the EcN-N12 treatment group had significantly lower TG and TC levels compared with the model group, the model + EcN group, the HFHCD group, the HFHCD + EcN group and the HFHCD + EcN-N12 group (Fig. 3d–g). Serum FFA levels were higher across all nicotine-treated groups, suggesting a link to nicotine exposure (Fig. 3h). Furthermore, ALT and AST activities were markedly reduced in the treatment group, indicating effective amelioration of liver injury (Fig. 3i–j).

**Fig. 3.**
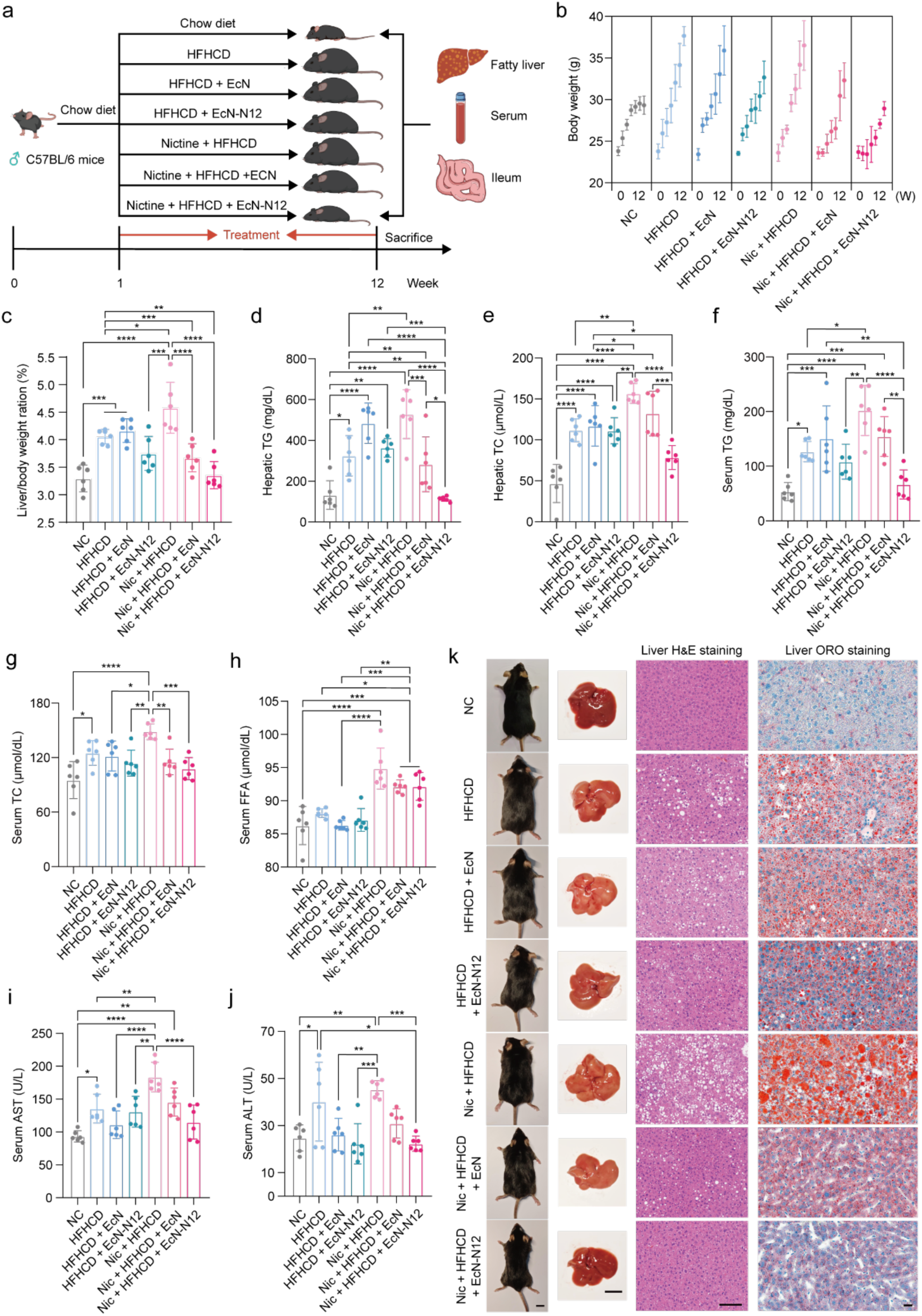
Evaluation of the therapeutic efficacy of EcN-N12 against nicotine-exacerbated NASH. a, Experimental scheme of EcN-N12 treatment in a mouse model of nicotine-exacerbated NASH. b, Changes in body weight of mice in each group (*n* = 6). c, Lung weight/body weight ratio of mice in each group (*n* = 6). d–e, TC and TG contents in the lung tissue of mice in each group (*n* = 6). f–h, Serum levels of TC, TG, and FFA in each group (*n* = 6). i–j, Serum AST and ALT levels in each group (*n* = 6). k, Representative photographs of body size, images of lungs, and hematoxylin–eosin (H&E) and Oil Red O staining of liver tissue sections from mice in each group (from left to right). Scale bars, 1 cm (body photographs), 1 cm (lung images), 200 μm (H&E), and 25 μm (Oil Red O). Statistical analysis was performed using one-way ANOVA with Tukey’s multiple comparison test. \**P* < 0.05, \*\**P* < 0.01, \*\*\**P* < 0.001, \*\*\*\**P* < 0.0001.

Morphological and pathological analyses further supported these findings (Fig. 3k). The EcN-N12 treatment group exhibited relatively lean body conformation, with livers retaining a healthy dark red color, in stark contrast to the greasy yellow appearance of livers from the model group. H&E staining revealed that nicotine promoted vacuolation resulting from hepatic lipid accumulation, a phenotype substantially mitigated in the EcN-N12-treated mice. Consistently, Oil Red O staining showed that nicotine increased both the density and size of lipid droplets in fatty livers, whereas only sparse lipid droplets were observed in the EcN-N12 treatment group. Collectively, these findings demonstrate that the engineered strain EcN-N12 effectively ameliorates nicotine-exacerbated NASH in mice.

## Discussion

This study systematically identified nicotine transporters and combined them with a key dehydrogenase (PpNicA^A107R^) and an electron carrier (CycN) to build a highly efficient nicotine-conversion module. Engineered probiotics carrying this module effectively cleared accumulated nicotine in the mouse gut and markedly inhibited the progression of non-alcoholic steatohepatitis via the gut-liver axis. These results, validated from engineered strains to animal models, establish the feasibility of probiotic-mediated harmless nicotine detoxification in the intestine and provide a promising translatable approach for combating nicotine-associated metabolic diseases. Native nicotine-metabolizing bacteria such as *P. putida* are limited by biosafety concerns due to their opportunistic pathogenicity^17^, while a recently identified gut nicotine degrader, *B. xylanisolvens*^9^, is hampered by its strict anaerobiosis and genetic intractability. By contrast, the probiotic EcN stands out as an ideal chassis, combining clinical safety with genetic amenability. Its therapeutic potential has been validated in ulcerative colitis management^45^, and more impressively, in the engineered strain SYNB1618 for phenylalanine metabolism in phenylketonuria^46^, underscoring its considerable potential for therapeutic metabolic engineering.

The pyrrolidine pathway employed in this study converts nicotine to *N*-methylmyosmine, a non-addictive intermediate that undergoes spontaneous hydrolysis to pseudooxynicotine. This is subsequently metabolized through a cascade of downstream enzymatic steps to 3-succinoylpyridine, 6-hydroxy-3-succinoylpyridine, and 2,5-dihydroxypyridine, ultimately yielding fumaric acid—which then enters the tricarboxylic acid (TCA) cycle for complete mineralization^32^. Beyond establishing a safe and nicotine-catabolic pathway, we identified and characterized high-performance nicotine transport elements, thereby addressing cellular uptake as a critical bottleneck limiting overall metabolic flux. Previous studies have suggested that bacteria are capable of saturable nicotine uptake^47^, but the underlying transporters have yet to be identified. Here, we identified two proteins, T7 (PPS_3866) and T3 (PPS_4370), that promote nicotine metabolism in EcN. T7 is a TolC-family outer membrane channel protein associated with the bacterial secretion and efflux machinery, whereas T3 is a hypothetical protein transcriptionally upregulated by nicotine in P*. putida* S16. Although neither protein is definitively established as the primary nicotine importer, our results (Figure 1d) strongly support their involvement in nicotine transmembrane transport. Elucidating their precise physiological functions and transport mechanisms will require further biochemical, structural, and genetic studies.

While these findings establish EcN-N12 as an effective hepatoprotective agent through gut nicotine clearance, several limitations merit consideration. Phenotypically, our analysis was largely confined to the gut-liver axis, leaving the systemic effects of nicotine detoxification on the brain, lungs, and vasculature to be determined. Translationally, the observed efficacy in mice, given interspecies differences in microbial ecology, nicotine handling, and MASH progression, requires confirmation in more clinically predictive models, and the long-term safety, stability, and ecological consequences of the engineered probiotic must be rigorously evaluated prior to therapeutic use. Addressing these questions will be a prerequisite for realizing the full translational potential of this approach.

In summary, this study developed a safe, efficient, and clinically translatable nicotine-degrading engineered probiotic, EcN-N12. This engineered strain can safely and efficiently eliminate intestinal nicotine, block gut-liver axis pathological signaling, and significantly alleviate nicotine-exacerbated NASH, offering a novel live biotherapeutic solution for nicotine-related metabolic diseases. Future research will focus on long-term safety evaluation, multi-model efficacy validation, in-depth mechanistic dissection, process optimization, and clinical trials to propel this engineered probiotic from the laboratory to clinical application, providing a novel armamentarium for the prevention and control of diseases associated with smoking and nicotine exposure.

## Method

### Strains and reagents

DH5α competent cells were purchased from TOLOBIO (Shanghai, China). EcN (*Escherichia coli* Nissle 1917) was obtained from the laboratory strain collection. LB medium: 5 g/L yeast extract, 10 g/L peptone, 10 g/L sodium chloride, dissolved in double-distilled water, and autoclaved at 121°C for 20 minutes. KOD DNA polymerase (Toyobo, Osaka, Japan) was used for high-fidelity amplification of DNA fragments. Recombinant plasmids were constructed using the ClonExpress Multis One Step Cloning Kit (Vazyme, Nanjing, China). The plasmid extraction and PCR product recovery kits were from (TransGen Biotech, Beijing, China). Primer synthesis, gene synthesis, and sequencing were all performed by Sangon Biotech (Shanghai, China). Arabinose, sodium salicylate, FAD, and FMN were purchased from Sangon Biotech (Shanghai, China).

### Plasmid construction

Taking the pUC19-P*_BAD_*-PpnicA-cycN plasmid as an example: using the P*_BAD_* plasmid as a template, the arac element and corresponding promoter were obtained via PCR using *arac*-F/*arac*-R primers; using the synthesized *PpnicA* as a template and the *PpnicA*-F/*PpnicA*-R primers, the *PpnicA* fragment was obtained via PCR; Using P*_tac_*-F and P*_tac_*-R primers, the P*_tac_* promoter fragment was obtained via template-independent PCR; using *cycN*-F/*cycN*-R primers with synthesized *cycN* as a template, the *cycN* fragment was obtained via PCR; using the pUC19 plasmid as a template, the linearized vector was obtained via PCR using pUC-bone-F/pUC-bone-R primers. The above fragments were assembled using the ClonExpress Multis One Step Cloning Kit (Vazyme, Nanjing, China), spread onto Amp-resistant LB plates, and incubated overnight at 37°C. Positive clones were identified via colony PCR and sequencing. Other plasmid construction methods followed a similar procedure; the specific primer sequences used are shown in Table 3.

### Constructing EcN genome-integrated exogenous genes

Single-copy integration: Taking the integration of the P*_BAD_*-PpnicA-cycN fragment at *ldhA* site as an example: First, the pCas plasmid was electroporated into EcN cells to obtain the EcN/pCas strain; the pTarget-*ldhA*-HA plasmid was electroporated into the EcN/pCas strain using the same method. After colonies grew, the integration of the target fragment was verified by colony PCR.

Using the same method, the P*_sal_*-TolC-H1 transporter expression cassette was integrated at the rhtB locus, ultimately yielding the EcN-N2 strain. The specific gRNA sequences are shown in Table 2.

Multi-copy integration: Using *Cargo*-F/*Cargo*-R primers with the pTarget-*ldhA*-HA plasmid as a template, P*_BAD_*-*PpnicA*-*cycN* expression cassette fragment was amplified by PCR. Using pCargo-F/pCargo-R primers with the pCargo plasmid as a template, a linearized vector was obtained by PCR. The above fragment and the linearized vector were then assembled into the pCargo-Nic plasmid using a homologous recombination kit; The pCAST-Array2 plasmid was obtained using the same method; pCargo-*nic* plasmid was electroporated into the 1×Nic strain and spread onto a Cm-containing LB plate; after obtaining positive clones, the pCAST-Array2 plasmid was electroporated into the corresponding positive clones using the same method and spread onto a Cm&Kan double-antibiotic LB plate; After obtaining strains containing the dual plasmids, dehydrotetracycline was used to induce expression of the array sequences and related elements on the corresponding vectors. The integration of the target fragments was verified via colony PCR, and the positive clone was named EcN-N4.

Using the same method, the engineered strains EcN-N6, EcN-N8, EcN-N10, and EcN-N12 were subsequently obtained. The specific crRNA sequences are shown in Table 2.

### Preparation of EcN competent cells and electroporation

Strip EcN onto an antibiotic-free LB agar plate and incubate overnight at 37°C. Pick a single EcN colony from the LB agar plate and inoculate it into 5 mL of antibiotic-free LB liquid medium. Incubate overnight at 37°C, 220 rpm, overnight. Subsequently, 300 μL of the bacterial culture was inoculated into 30 mL of LB liquid medium and cultured until the OD_600_ reached approximately 0.4. The cells were then washed three times with 20 mL, 10 mL, and 5 mL of 10% glycerol (4°C), and finally resuspend in 300 μL of 10% glycerol, then aliquot 100 μL per tube into 1.5 mL centrifuge tubes.

Using the pUC19-P*_BAD_*-*PpnicA*-*cycN* plasmid as an example, transform 0.2 μg of the plasmid into EcN electroporation-competent cells. After electroporation, resuspend the cells in 1 mL SOC and incubate with shaking (220 rpm) at 37°C for two hours. Subsequently, plate all cells onto LB agar plates containing 100 μg/mL ampicillin, incubate overnight at 37°C, and confirm the successful transformation of the target vector into EcN via colony PCR the following day.

### Induction culture of EcN-N12

Inoculate EcN-N12 into 5 mL of LB liquid medium and incubate overnight at 37°C and 220 rpm to obtain a primary seed culture. The next day, transfer 5% of the cultured bacterial suspension to 30 mL of LB liquid medium and continue culturing at 37°C and 220 rpm until the OD_600_ reaches 0.8. Add 1 M arabinose solution and 100 mM salicylic acid solution to the engineered bacterial culture to induce expression for 3 h. Centrifuge at 6000 rpm for 10 min, and collect the bacterial cells at an OD_600_ of 5. Resuspend in PBS buffer, centrifuge, and repeat this process twice before resuspending the cells in PBS buffer once more.

### HPLC analysis

Incubate the induced EcN-N12 suspension with a 100 μg/mL nicotine solution at 37°C and 220 rpm for 2 h. Centrifuge the solution at 12,000 rpm for 10 min, transfer the supernatant to a new Eppendorf tube, and filter through a hydrophilic membrane for later use. Determine the nicotine concentration in the supernatant using a high-performance liquid chromatograph (Agilent, USA).

Substrate Concentration Assay: The catalytic activity of EcN-N12 was assessed by measuring the residual nicotine using HPLC under the following conditions: Column: C18 column, 4.6 × 250 mm, 5 μm, 120A; Mobile phase: KH_2_PO_4_ (10 mM): methanol: trimethylamine = 90:10:0.1 (V: V: V); Column temperature: 30℃; Flow rate: 0.6 mL/min.

Calculation of results: First, peak areas were converted to corresponding concentration values using a standard curve. The concentration of the blank control (without added bacteria) was taken as the initial concentration, and the concentration of the sample detected by HPLC was taken as the residual concentration. The difference between the two represents the amount of nicotine consumed by the EcN-N12 strain.

### Determination of nicotine-degrading ability of EcN-N12 in a simulated intestinal Environment

Nicotine (Sigma, #54-11-5) was added to artificial small intestinal fluid and artificial colonic fluid (Aladdin, Shanghai, China) to achieve a final concentration of 20 μg/mL, simulating the environment of nicotine accumulation in the human intestine. A suspension of induced EcN-N12 cells was thoroughly mixed with artificial small intestinal fluid and artificial colonic fluid containing 20 μg/mL nicotine, respectively. The mixtures were incubated at 37°C and 220 rpm for 2 h. After incubation, the residual nicotine content in the system was determined using HPLC.

### Animals

C57BL/6 mice were purchased from Shanghai BK/KY Biotechnology Co., Ltd. The mice were male, 8 weeks old, weighed approximately 20 g, and were fed SFP-grade chow. The ethics approval number was IRB-AF63-V1.0.

### Mouse model of nicotine-associated NASH

Forty-two C57BL/6 mice were randomly divided into 7 groups (*n*=6) and fed a normal diet with normal drinking water for one week. Starting in the second week, one group was fed a normal diet and normal water, three groups were fed a high-fructose and high-cholesterol diet (Trophic, #TP26304, Nantong, China) and normal water, three groups were fed a high-fructose and high-cholesterol diet (HFHCD) and water containing nicotine (0.2 g/mL, Sigma, #54-11-5) in “sweet water” containing 23.1 g/L D-(−)-fructose (Sigma, #F0127) and 18.6 g/L D-(+)-glucose (Sigma, #G8270)^9, 48^. Following the above protocol, seven groups of C57BL/6 mice were fed continuously for three months. During this period, the three groups of mice on HFHCD were gavaged every two days with PBS buffer, EcN, and EcN-N12, respectively, while the three groups of mice on the 0.2 g/L nicotine-sweetened water diet were gavaged with PBS buffer, EcN, and EcN-N12, respectively. After three months, mice in each group were euthanized. Liver tissues were photographed, weighed, and subjected to H&E and Oil Red O histological staining (RecordBio, Shanghai, China). Biochemical parameters including TG and TC were measured in the liver (Boxbio Science & Technology Co.,Ltd., Beijing, China). Tissues from the heart, spleen, lungs, and kidneys were photographed and recorded, and these organs were also subjected to H&E histological staining. Liver, heart, spleen, lung, and kidney tissues were ground and spread on slides to determine the bacterial load in each organ. Serum samples were collected from mice in each group to measure biochemical parameters such as TG, TC, FFA, AST, and ALT (Boxbio Science & Technology Co.,Ltd., Beijing, China), as well as inflammatory markers including IL-6 (MULTI SCIENCES, #EK206) and TNF-α (MULTI SCIENCES, #EK282).

## Notes

### Competing Interest Statement

The authors have declared no competing interest.

